# Ecological change points: The strength of density dependence and the loss of history

**DOI:** 10.1101/224352

**Authors:** José M. Ponciano, Mark L. Taper, Brian Dennis

**Affiliations:** Department of Biology, University of Florida, Gainesville, FL. 32611.; Department of Ecology, Montana State University, Bozeman, MT, 59717; Department of Fish and Wildlife Sciences and Department of Statistical Science, University of Idaho, Moscow ID 83844-1136, USA

**Keywords:** Strength of density dependence, Gompertz model, breakpoint, change-point stochastic processes, state-space models

## Abstract

Change points in the dynamics of animal abundances have extensively been recorded in historical time series records. Little attention has been paid to the theoretical dynamic consequences of such change-points. Here we propose a change-point model of stochastic population dynamics. This investigation embodies a shift of attention from the problem of detecting when a change will occur, to another non-trivial puzzle: using ecological theory to understand and predict the post-breakpoint behavior of the population dynamics. The proposed model and the explicit expressions derived here predict and quantify how density dependence modulates the influence of the pre-breakpoint parameters into the post-breakpoint dynamics. Time series transitioning from one stationary distribution to another contain information about where the process was before the change-point, where is it heading and how long it will take to transition, and here this information is explicitly stated. Importantly, our results provide a direct connection of the strength of density dependence with theoretical properties of dynamic systems, such as the concept of resilience. Finally, we illustrate how to harness such information through maximum likelihood estimation for state-space models, and test the model robustness to widely different forms of compensatory dynamics. The model can be used to estimate important quantities in the theory and practice of population recovery.

## 1. Introduction

Change more than stasis has perplexed both theoretical and empirical students of ecological time-series. Questions like: “when is a change in dynamics going to occur?”, or “why does a change in dynamics occur?” have preoccupied ecologists working in population dynamics for decades now [1]. Yet, relatively missing in the theoretical population 5 ecology literature (but see [2]) is an in-depth exploration of how a change in a ecological process, such as density dependence, can drive change in the statistical properties of a (inherently stochastic) dynamic system. How these in turn result in changes with direct management implications is a question of paramount relevance in conservation biology.

The origin of this paper can be traced to a theoretical and empirical study of the population dynamics of a salmonid species by members of the Taper laboratory and colleagues [3, 4, 5, 6, 7, 8, 9]. In particular, we were then aiming at modeling the reaction of the population dynamics of a Bull Trout population (Salvelinus confluentus) to a drastic change in the community composition. Such change was noticeable by eye in the trends of the annual counts of adult females (data from the Montana Fish and Wildlife Service): around 1991, a marked drop in abundances consistently occurred in different tributaries of the affected river and lake system. This scenario led us to specify a natural model candidate: a population dynamics model with a change-point. After all, that very same statistical model had long been used in statistical time series modeling. Managers were and are eager for answers to questions like: how were the dynamics before the change affecting the dynamics after the change, and the extent of the change? After so many years fluctuating at low abundances, is the population expected to recover? If so, how long does the recovery process would take?

The strength and effect of density dependence has long been a focus of theoretical and applied population ecology [10, 11]. Although the dependence of the per capita growth rate of a species on its own density or abundance is now widely regarded as a main driver of population dynamics, such proposition still instills rich debates, theoretical problems and practical dilemmas (see citations in [12], [13], [14]). Our approach directly links a change in dynamics with this central concept in theoretical ecology.

The purpose of this paper is to show that progress in understanding population trends with a change point does not reside per se in the implementation of statistical methodologies aiming at detecting and quantifying change. Rather (through the analysis of these Bull Trout population trends) we found that hidden within the numerical evaluation of the standard matrix model specification of the change-point auto-regressive moving average model, was an ecologically meaningful mechanistic process. Clear patterns and coherent algebraic structures appeared that showed how the strength of density dependence shapes the post change-point dynamics. These expressions partitioned the variance components of the stochastic population process in a way that it allowed us to directly answer those important managerial questions through changes in the strength of density-dependence. Surprisingly then, parameter estimation turned out to be a simple by-product of the process of scientific inquiry of what a stochastic model was really implying.

We strongly believe that the results presented in this paper are generally important for two reasons: first, these results have been all along within the reach of any statistical analysis of population time series data but have so far been overlooked, hidden within the associated matrix algebra calculations of any statistical time series analysis. Second and most importantly, to our knowledge no other work extracts from routine matrix algebra calculations like we do, the explicit and readily interpretable algebraic forms that allow the analyst to go back and forth between the statistical and numerical phrasing of a change and its ecological interpretation.

Statistical time series models with change points in population dynamics have long been used [1] but are nowadays rarely routinely considered in ecological analyses despite being extensively studied in the statistical literature (but see [15, 16, 17, 18, 19]). Asymptotics of maximum likelihood parameter estimates have been derived [20, 21, 22, 23]. Hypotheses tests for the presence of change points have been proposed [24, 25, 26, 27, 28, 29]. Quality control theory features studies of how to incorporate change point warnings in control charts [30, 31, 32]. These statistical methods usually test whether an observed time series of population abundances (or densities) depart from a fixed deterministic equilibrium, or from a stochastic stationary distribution. In the first approach, the observed population abundances are observations with measurement error (observation error) added to a deterministic equilibrium or trend. In the second approach, stochastic perturbations are included in the dynamic model to describe the natural fluctuations in abundance. These natural fluctuations are also known as “process noise” [7]. With process noise, concepts from deterministic ecological accounts of population regulation take on new forms. For instance, a stable equilibrium of population abundances becomes a stationary probability distribution in stochastic population models [33]. Although these statistical studies provide substantial underutilized opportunities for ecological data analysis, missing to date is an ecological understanding of the dynamical properties of populations undergoing a change point.

The last twenty years of research in statistical population dynamics concentrated on parameter estimation, model selection and hypothesis testing of models with and without process and observation error [12, 34, 35, 36, 37, 7, 38, 39, 40, 41, 42, 43]. This research program is warranted because the effective statistical coupling of empirical observations with mathematical models is critical for testing hypotheses concerning population regulation [44]. However, the ecological investigation of populations undergoing change has lagged behind the attention to statistical issues [7, 45]. This paper seeks to understand what are the ecological consequences of population processes that are not static, but undergo change. To do that, we reveal how the analytical phrasing of a change in a dynamic model translates into readily interpretable and closed-form mathematical changes in the stochastic properties of the population process.

The key idea of this paper is to represent an ecological change point as a saltational change in the parameters of the stochastic population model. Such a change produces both a shift in the stationary distribution of population abundance and a transition to a new equilibrium distribution. We obtain simple formulae characterizing the statistical distribution of the population process during the transition between stationary distributions of abundance. Our study demonstrates that certain historical properties of a population’s growth dynamics are determined by the strength of density-dependent processes. In particular, a time series transitioning from one stationary distribution to another contains information about where the process was before, where is it heading, and how long it is going to take to get there. We illustrate how to harness the information in time series containing transitionary portions via maximum likelihood estimation for state-space models. The model proposed here provides new insights into the role of density dependence in shifting environments. This model also represents a practical tool for detecting and predicting change events in population monitoring.

## 2. The strength of density dependence in discrete time models

Consider the general discrete time population growth model *n*_*t*+1_ = *f*(*n_t_*) = λ(*n_t_*)*n_t_*, where *n_t_* is the population density or abundance at time *t* and λ(*n_t_*) is the (density-dependent) per capita growth rate. Assume that *f*(*n*), the recruitment map, is continuously differentiable and that *n\** is its non-trivial equilibrium abundance (*i.e.* its satisfies f (*n\**) = *n\**).

Three measures of the strength of density dependence have been suggested. The first measure is motivated by thinking of the strength of density dependence simply as the marginal effect on the per capita growth rate of an increase in density [10, 46], which according to our general setting, corresponds to *∂*λ(*n_t_*)/*∂n_t_*. This is a measure of the effects of density at the individual-not population-level that has been used, among other things, to phrase an evolutionary perspective of intra-specific competition as the way individual, demographic traits respond to an increase in density [46]. Because for some discrete, density-dependent maps the per capita growth rate λ(*n_t_*) is written as an exponential function (as in the Ricker or in the Gompertz equations, see [12, 11]), the marginal effect of an increase in density can be conveniently measured and plotted in the log-scale of the per capita growth rate by computing *∂* ln λ(*n_t_*)/*∂n_t_*.

The second measure corresponds to the derivative of the recruitment map *f*(*n*) at equilibrium. Such measure can be directly read from the graph of *f*(*n*) as a function of n as the value of the slope when *f*(*n*) crosses the 1:1 line. It is important to note that a consideration of both an individual and a population wide measure of density dependence is of obvious importance in population ecology to explain persistence, average abundance, bounds on temporal stability and in general, for a careful understanding of the unfolding of population dynamics [46].

A third measure of the strength of density dependence in a simple model with no age structure was given by Lande et al (2002), who defined it as the negative elasticity at equilibrium of the per capita population growth rate with respect to change in the population. This amounts to computing 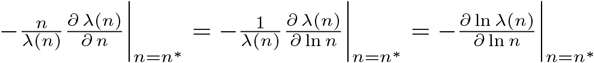.

This definition of the strength of density-dependence is insightful and relevant in population dynamics modeling, not only because it provides a scale-free measure of the change in the per capita growth rate as the (log) population size varies, but also because it is readily extendable to scenarios dealing with more complex life histories [47].

## 3. Methods

### 3.1. The discrete Gompertz model

The deterministic skeleton of our stochastic model is the discrete-time Gompertz model [7, 11] given by:

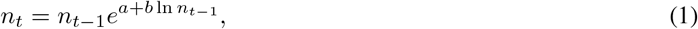

where *n_t_* is population abundance at time *t* and *a* and *b* are constants. Note that many formulations of the Gompertz model explicitly include a minus sign before b and thus can look superficially different.

The non-trivial equilibrium density, or carrying capacity is given by *n\** = *e^−a/b^*. This equilibrium is asymptotically stable, provided the absolute value of the derivative of the recruitment map (eq. 1) evaluated at *n\** is less than 1 [48], i.e., if |1 + b| < 1. For this model, the strength of density dependence measured using the per capita growth rate is *∂* λ(*n_t_*)/*∂n_t_* = *be*^*a+bln n_t_*-1^ /*n_t_* which is equal to *b*/ exp{-*a/b*} at equilibrium. The measure of the strength of density dependence using the logarithm of the per capita growth rate, written as *∂*ln λ(*n_t_*)/*∂n_t_* = *b/n_t_* is again equal to *b*/ exp{‒*a/b*} at equilibrium. Finally, Lande et al.’s negative elasticity is simply given by −*b* for all population abundances. Then, the density dependent coefficient *b*, or its carrying capacity-scaled version *b*/ exp{−*a/b*}, is directly involved in the definition of the strength of density dependence. So following [47, 49, 11], heretofore we identify *b* as the strength of density dependence for the Gompertz model, due to its role in the rate of approach to a locally stable equilibrium.

Let *x_t_* = ln(*n_t_*) and *c* = *b* + 1. Then, Equation (1) becomes the first order difference equation:

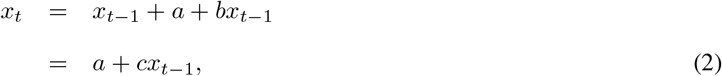

whose solution can be found by induction

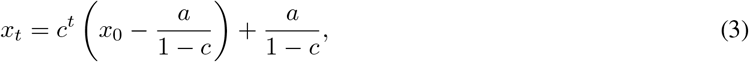

provided *c* ≠ 1. If *c* = 1 then *b* = 0 and population growth is density independent. The solution in that case is *x_t_* = *x*_0_ + *at*. If density-dependent processes lead to a stable equilibrium density, *i.e,* if |c| < 1 then we may distinguish two cases: 0 < *c* < +1 and −1 < *c* < 0. In both cases, as *t* → ∞, the log population density (abundance) 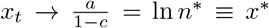. In the first case, *x_t_* approaches *x\** directly, either from above or below, depending on initialization. In the second case, the deterministic map exhibits damped oscillatory behavior while approaching the equilibrium.

### 3.2. The stochastic Gompertz model with environmental noise

A model of Gompertz population growth with environmental stochasticity may be written as

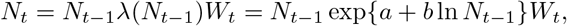

where the population size *N_t_* is now modeled with a random variable representing environmental variation. A useful model for the noise takes the *W_t_* to be independent and identically distributed (*iid*) log-normal random variables describing heavy-tailed temporal perturbations of the per-capita growth rate [50]. Accordingly, we set *W_t_* = *e^E_t_^* where *E_t_* ~ N(0, *σ*^2^) to get

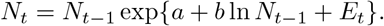

On the logarithmic scale, with *X_t_* = ln *N_t_*, this stochastic Gompertz model with environmental noise becomes the well known AR(1) process

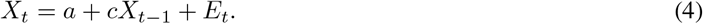

Provided |*c*| < 1, this stochastic process has a normal stationary distribution with mean and variance given by

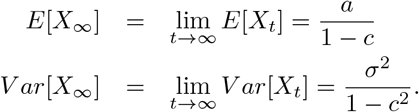

Thus, both, the process stationary mean (equivalent to the deterministic stable equilibrium) and its associated variance critically depend on the strength of density dependence. Simply put, although external, environmental variability induces a variability in the growth rate, the magnitude and intensity of the population fluctuations are also influenced by the strength of density dependence. This feature led Ives et al (2003) to a detailed study of the concept of ecological stability applied to stochastic community dynamics models. Before moving on to the breakpoint model, we note that through this article, random variables will denoted with capital letters and realizations of these random variables with lower case letters. Also, random vectors like the multivariate random vector ***X*** = [*X*_0_, *X*_1;_ *X*_2_,…, *X_q_*]′, where the *X_i_* denote the log-population abundance at time *i*, will be denoted with capital, boldface letters. Realizations of the random vectors will be written with lower case, bold face letters. Finally note that here, as in all the vectors we define, the length of the vector is the last index (*q* in this case) plus one.

### 3.3. The breakpoint Gompertz model

We consider the case where the stochastic process model undergoes a change after a given time time step, *τ*. Operationally, such change is implemented by assuming that the value of one or more of the stochastic Gompertz model parameters changes after time *τ*. A general model, from which all of the possible changes can be specified as sub-models, specifies that there are two different sets of parameters for the stochastic Gompertz model before and after the change-point. If *Z_t_* denote *iid* N(0,1) random variables, a general change-point model is written as

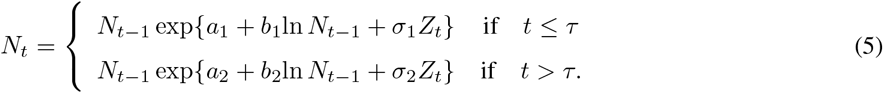

In the log-scale, the model becomes

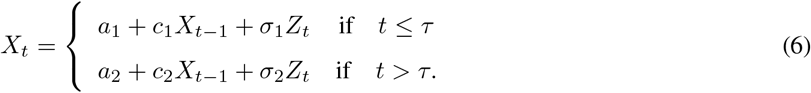

Here, the environmental noise process is written as *σ_i_Z_t_* to show model parameter changes after the breakpoint, as opposed to *E_t_* as above. Also, whether or not *t* < *τ* will determine if 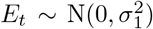 or 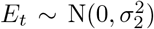, so no additional subscript or notation will be used from now on to distinguish between the environmental noise random variables before and after the breakpoint. Also, as before, we assume that |*c*_2_| < 1. Writing the breakpoint model as in equation 6 is a common practice, for instance, among studies of applications of Random Markov Fields in Biology [51]. However, as we show below, writing the breakpoint model in such form, and/or its equivalent matrix representation falls short of a full analysis of the role of the strength of density dependence in shaping the speed of the change and all statistical properties (means, variances and covariances) of the process. After writing the model in its matrix format, we did the explicit matrix algebra multiplications, simplified the expressions and sought out to write the moments of the process as a function of the strength of density dependence. The resulting formulae as well as their ecological interpretation are the main results of this paper. To begin these calculations, by induction (see Appendix 1), we arrive at a matrix equation for the entire random vector of the log abundances before and after the breakpoint:

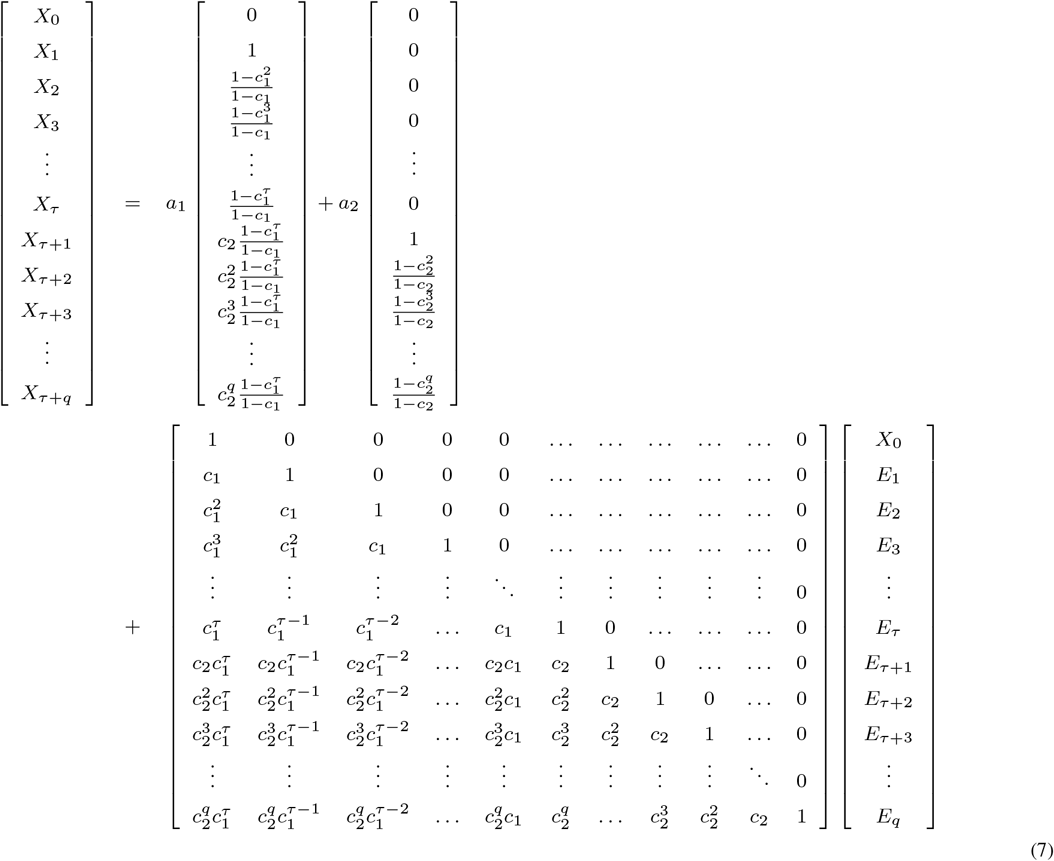

or

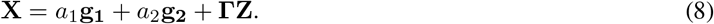

Note that vector **Z** is Multivariate Normal (*MVN*) with mean

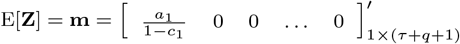

and variance-covariance matrix

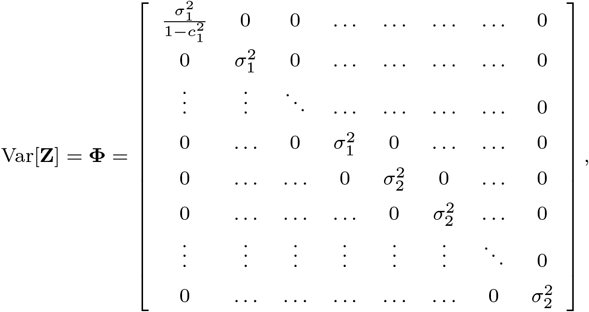

where the changes in the diagonal occur in the 2^nd^ and in the (*τ* + 2)^th^ elements. Because a linear transformation of a multivariate normal distribution is again multivariate normal (Theorem 3.3.5 p. 101 Graybill 1976), it immediately follows that

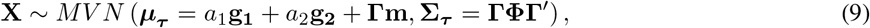

where the subscript *τ* indexes the mean and the variance of the process with a change-point, to explicitly differentiate them from the mean *μ* and variance *Σ* of the process without a change-point (see Appendix 1).

### 3.4. Parameter estimation

A state space model where sampling error is compounded to the process error can be easily specified, and that is useful for parameter estimation. We call this model the Break-Point Gompertz State-Space (BPGSS) model. Using equation 9 and a lognormal sampling error process (see the discussion of [7] for a justification of such model), the multivariate distribution of the log-observations with added sampling error **Y = X + F** is

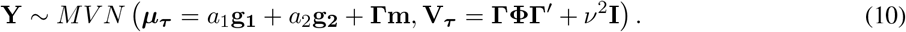

This *MVN* distribution for the observations follows since both, the pdf for the process *X* and for the observation error *F* are both *MVN*, just as in the model of [7]. Parameter estimation via Maximum Likelihood (ML) for this state space model then proceeds by maximizing this multivariate pdf over the model parameters. This pdf can be readily programmed as it is written above, but the computations can be rendered much more efficient when the matrix multiplications in expression (10) are carefully simplified (see Appendices 1 and 2). Sample code for parameter estimation is uploaded as supplementary material and available upon request from JMP.

### 3.5. Testing the robustness of the BPGSS model

Clearly, the BPGSS model is a simplified representation of a population dynamics undergoing a drastic change. One of the most important questions for practicing ecologists is to what extent these simple population dynamics models can be used to infer intrinsic properties of the system that can be of immediate utility. Therefore, we performed a validation effort via simulations that aimed at testing the robustness of the quantitative and qualitative model inferences derived from this model. To do that, we simulated breakpoint dynamics by relaxing most of the major simplifying assumptions made by the BPGSS model and adding biological complexity. Then, the simulated dynamics *(i.e.* time series) were saved and used to confront the BPGSS model with different ecological scenarios of interest.

The discrete time, discrete state stochastic process used to simulate population dynamics was a Markov process that, besides incorporating sampling error, accounted for demographic stochasticity and different forms of environmental variability. Most of the environmental noise models, like the BPGSS model, incorporate stochasticity in the maximum growth rate. Such noise is multiplicative in the scale of population abundances, and additive in the scale of the logarithmic per capita growth rate. Another possibility, however, is to assume temporal randomness in the strength of density dependence. This approach assumes that the environmental forces shaping the population dynamics affect the intensity of density-dependence rather than the maximum growth rate and to date, remains a rarely explored model alternative (but see [52, 53]).

Our discrete-time Markov model of reproduction and survival followed the general discrete time model construction presented by [54, 55] and later revisited and expanded by [56] and [53]. A detailed explanation of such construction is given in the supplemental material of [56]. Briefly, population growth from one generation to the other is modeled as a two-stage, stochastic process: at time *t* the *n_t_* adults present in the population give birth each to a random number of offspring (*i.e.* 0,1,2,3, …). Then, we assume that both adults and offspring survive to time *t* + 1 according to a density dependent survival probability, which takes on one of the well-known discrete-time density-dependent maps (Ricker, Hassell, Gompertz, Below, etc…). This reproduction and survival process considers only demographic stochasticity. To include environmental stochasticity, [54] and [55] have previously assumed before that the mean of the offspring distribution varies randomly from one time step to the next according to a stochastic environmental process. Such variability mimics the random, temporal fluctuation in the quality of the environment that leads to temporal fluctuations of the average number of offspring produced by every adult. As recently shown by [53], this is but one way of incorporating environmental variability. Here, we assumed that the environment randomly perturbs both the mean of the offspring distribution and the density dependent survival probability. For our simulations, we assumed that the random (over time) environmental process, denoted Λ, had independent and identically distributed realizations from a Gamma(*k, α*) distribution. The results we present here use the value of the scale parameter α set to 1000 and the shape parameter *k* set to 1500, but hold for other parameter values as well.

Accordingly, conditional on the realized value λ of the random environmental process, each individual, independently from each other, was assumed to have a Poisson (λ) offspring distribution. As a result, from time *t* to time *t* +1, the (conditional) number of potential recruits was distributed Poisson (*n_t_*λ). We then modeled offspring survival with a density dependent, binomial thinning process. The survival probability of the offspring (potential recruits) was taken to be an explicit function of the population size (density) at time *t*, denoted as *n_t_* by writing it as

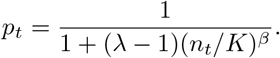

This function for the offspring survival probability *p_t_* corresponds to Below’s recruitment map [57]. The adoption of this function is important because it provides a means to test the generality of our results under different forms of density dependence. Indeed, in this function *p_t_* the parameter *β* controls the type of compensatory dynamics (under compensatory (*β* < 1), compensatory (*β* = 1) and over compensatory (*β* > 1). Thus, tuning the value of the parameter *β* effectively changes the shape and curvature of the recruitment map and can recapitulate the behavior of an entire variety of discrete recruitment maps (see [57] for details).

Accounting for binomial survival when the number of potential recruits is Poisson distributed is well known to be a special case of a “randomly stopped sum” and results in a Poisson distributed total number of recruits from one time to the next [54]. Here, the (random) total number of recruits N for the next generation, is denoted as N_t+1_. From the previous assumptions, it follows that N_t+1_|(N_t_ = n_t_, Λ = λ) ~ Poisson (λn_t_p_t_). Note again that because the realized value of the environmental process λ modifies the density dependent term in Below’s model, the effect of the stochastic environmental process we specified is qualitatively different from the environmental noise formulation proposed in our BPGSS model. This stochastic model formulation where the stochastic environmental process affects the density-dependent coefficient was recently studied by [53]. Introducing this type of variability changes the well-known scaling relation between the population variance and population abundances in stochastic growth models, and may be important in many animal populations [53]. Averaging the environmental process over time results in the marginal transition pdf of the process. This transition, however, does not result in the well-known demographic and environmental negative binomial process [54], because of the complex environmental process effects.

To incorporate different density-dependent scenarios, we used three values for *β*: 0.5, 1 and 1.2. These values represent respectively under-compensatory, compensatory and over-compensatory dynamics. This axis of variation allowed us to test the robustness to different forms of density dependence. The simulations were started at the value of the deterministic skeleton’s carrying capacity in the log-scale (14) and a change point of dynamics was introduced at time *t* = 40. The change consisted of a change in the deterministic skeleton carrying capacity, from 14 to 4.48 (in the log-scale). Although we introduced a single change at a single time point, for each value of *β* it took some time for the time series to “stabilize” around the second deterministic carrying capacity. The time it takes for the average of the process to reach the arithmetic average of the two (log) deterministic carrying capacities is heretofore referred to as the “half-life of the change”. The resulting simulated population dynamics was sampled 1000 times per value of *β* using a multiplicative lognormal sampling error with parameters μ = 0 and ν^2^ = 0.2315. This sampling resulted in 1000 time series of length 200 per value of *β* that were taken as data sets to which the BPGSS model was fitted. Furthermore, for each value of *β*, the “true” half-life of the change point was computed numerically as the average half-life in 50000 simulated time series. We then computed the relative bias of our half-life estimator for each one of the 1000 simulated time series to which the BPGSS model was fitted.

## 4. Results

### Understanding the breakpoint dynamics

The vector expressions in the multivariate normal distribution of the log-normal abundances (eq. 9) conceal explicit, simple expressions linking the dynamics before and after the change. In particular, immediately after the change, there is a transitional period. The time-varying distribution of this transition contains information about both, the past stationary mean and variance, as well as the future stationary mean and variance. This information is exposed by partitioning the vector X of log-population abundances over time, into a vector of pre-breakpoint log-abundances **X_1_** = [*X*_0_ *X*_1_ *X*_2_ … *X*_τ_]′ and another of post-breakpoint log-abundances **X_2_** = [*X*_*τ*+1_ *X*_*τ*+2_ … *X*_τ+q_]′. In what follows, we state six main results along with three corollaries that follow from this partitioned structure. These results follow from a detailed algebraic examination of the vector and matrix calculations resulting from the model formulation in equations 7-9. These algebraic examinations, necessary to fully understand the results, are presented in appendices 1 and 2. Each one of these results brings a different dimension to the understanding of the transitional dynamics in ecological terms. At the end, we also list the result regarding the robustness tests of the BPGSS model.

**Result 1:** *In transition, the mean of the (log) population process becomes a weighted average of the stationary mean before the breakpoint, and the stationary mean predicted by the new set of dynamical parameters. The weights of this weighted average are given by 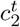 and 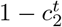 respectively, where t is the number of time steps after the break.*

The result follows directly from the expression for the mean of the multivariate distribution in eq. 9. By developing and explicitly writing the entries of the vector and matrix calculations in this expression, we found that the mean vector of the first partition **X_1_** is given by 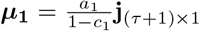, where **j**_(τ+1)×1_ is a vector of ones, whereas the mean of the second partition is given by the time-varying weighted average:

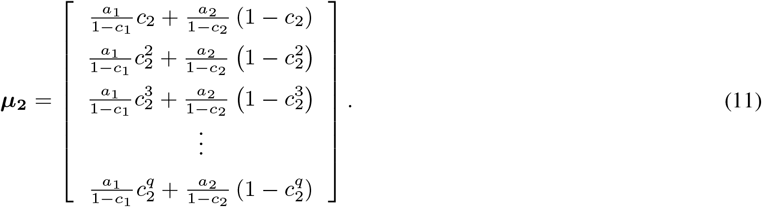

The first element in the vector above (eq. 11) is the mean of the transition process one time step after the breakpoint. This element is a weighted average of the first stationary mean *μ*_1_ = *a*_1_/(1 – *c*_1_) and the second stationary mean *μ*_2_ = *a*_2_/(1 – *c*_2_). The weights are given by the strength of density-dependence after the breakpoint, *c*_2_. Thus, this coefficient completely determines how important the mean of the first phase of the process is right after the change. Two time-steps after the breakpoint, the weights are given by 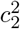 and 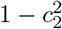 respectively. Again, because 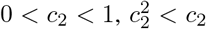 and hence, the influence of the first stationary means decays two time steps after the breakpoint. *t* time units after the breakpoint, the weights become 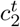 and 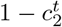. As a result, as the number of time steps *t* after the breakpoint grows large, the weight for the first stationary mean goes to 0 while the weight of the second stationary mean converges to 1. It then follows that the mean of the post-breakpoint process truly represents a transitionary trend.

**Result 2:** *The speed of the change of the mean of the log-population process (eq. 9) is determined by the strength of density dependence. The half life of the change in means, or the average time 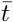 it takes for the average of the process to reach (*μ*_1_ + *μ*_2_)/2 from *μ*_1_, is a quantity that directly depends on the strength of density dependence post breakpoint. Thus, any property of this dynamic system that depends on the waiting time to complete the transition, also depends on the strength of density dependence.*

An explicit expression for the half life of the transition in means is readily found by setting

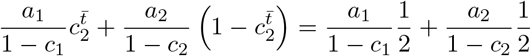

and solving for 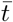, which yields

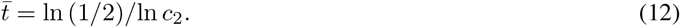

Thus, the speed of the change from one stationary mean to the other one is determined by the strength of density dependence after the breakpoint. For example, if *c*_2_ is close to 0, then the transition weights 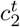 and 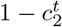 (see eq. 11) converge fast to 0 and 1, respectively and the process approaches the second stationary mean faster. A detailed characterization of the speed of the change as a function of *c*_2_ is given in Figs. 1 and 2. From this characterization it is readily apparent that when 1/2 < *c*_2_ < 1 the strength of density dependence can be thought as weak; strong when −1/2 < *c*_2_ < 1/2 and as very strong when −1 < *c*_2_ < −1/2 (see legend in Fig. 2).

**Figure 1:**
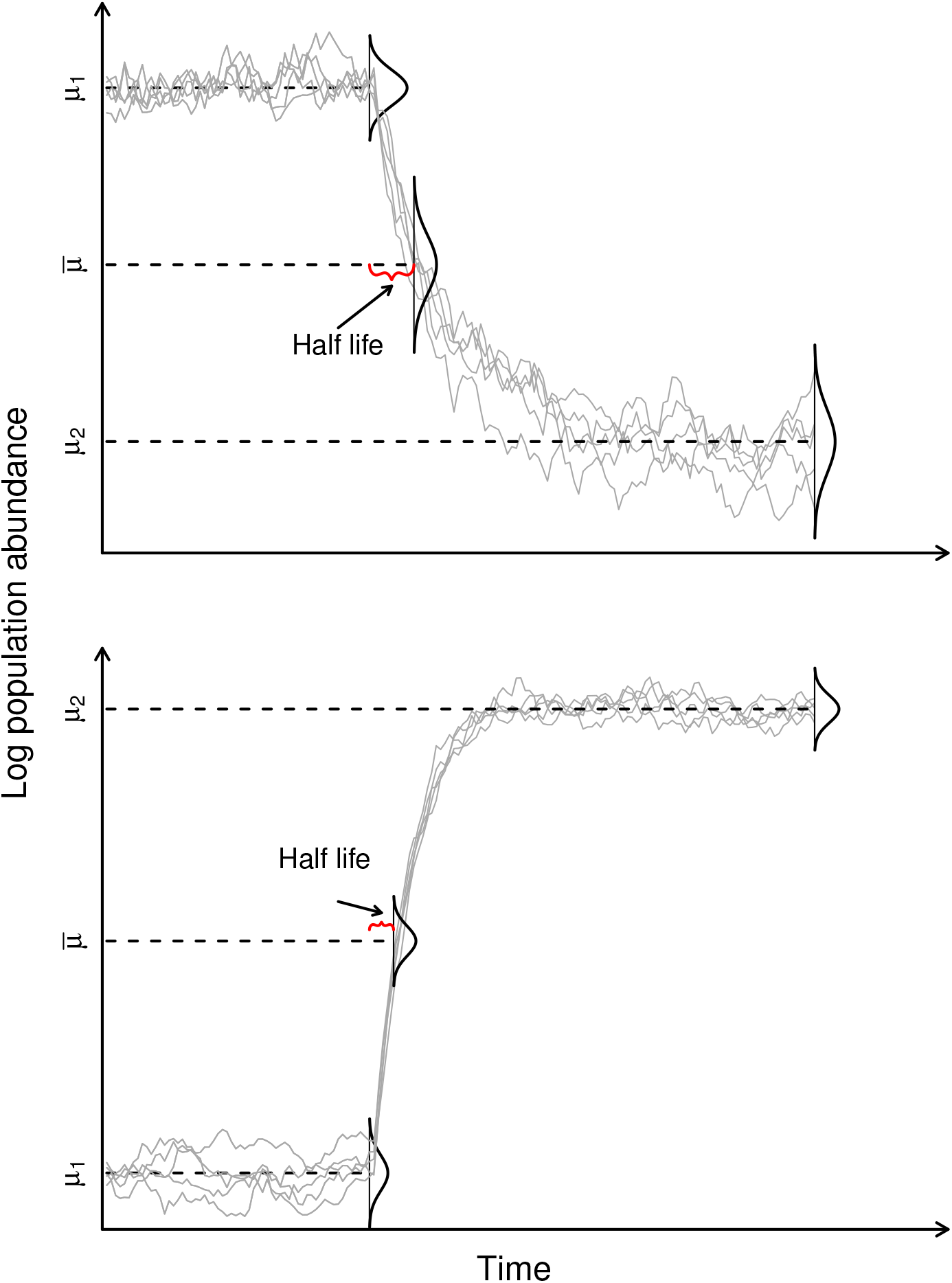
The stochastic Gompertz breakpoint process and the half life of the change in the mean of the dynamical process, 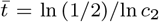. Plotted are 5 realizations of the Stochastic Gompertz breakpoint process. Dotted lines mark the process mean before the breakpoint (*μ*_1_ = *a*_1_/(1 – *c*_1_), the mean after the breakpoint (*μ*_2_ = *a*_2_/(1 – *c*_2_), and the arithmetic average of both means 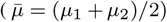. The time at which such arithmetic average is reached is the half life of the process, 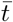. The upper panel shows a case where the mean changes to a smaller size whereas the lower panel depicts the change from a lower equilibrium size to a higher one. This second case could depict a population recovery scenario, in which case the half life is half the mean time to recovery.

**Figure 2:**
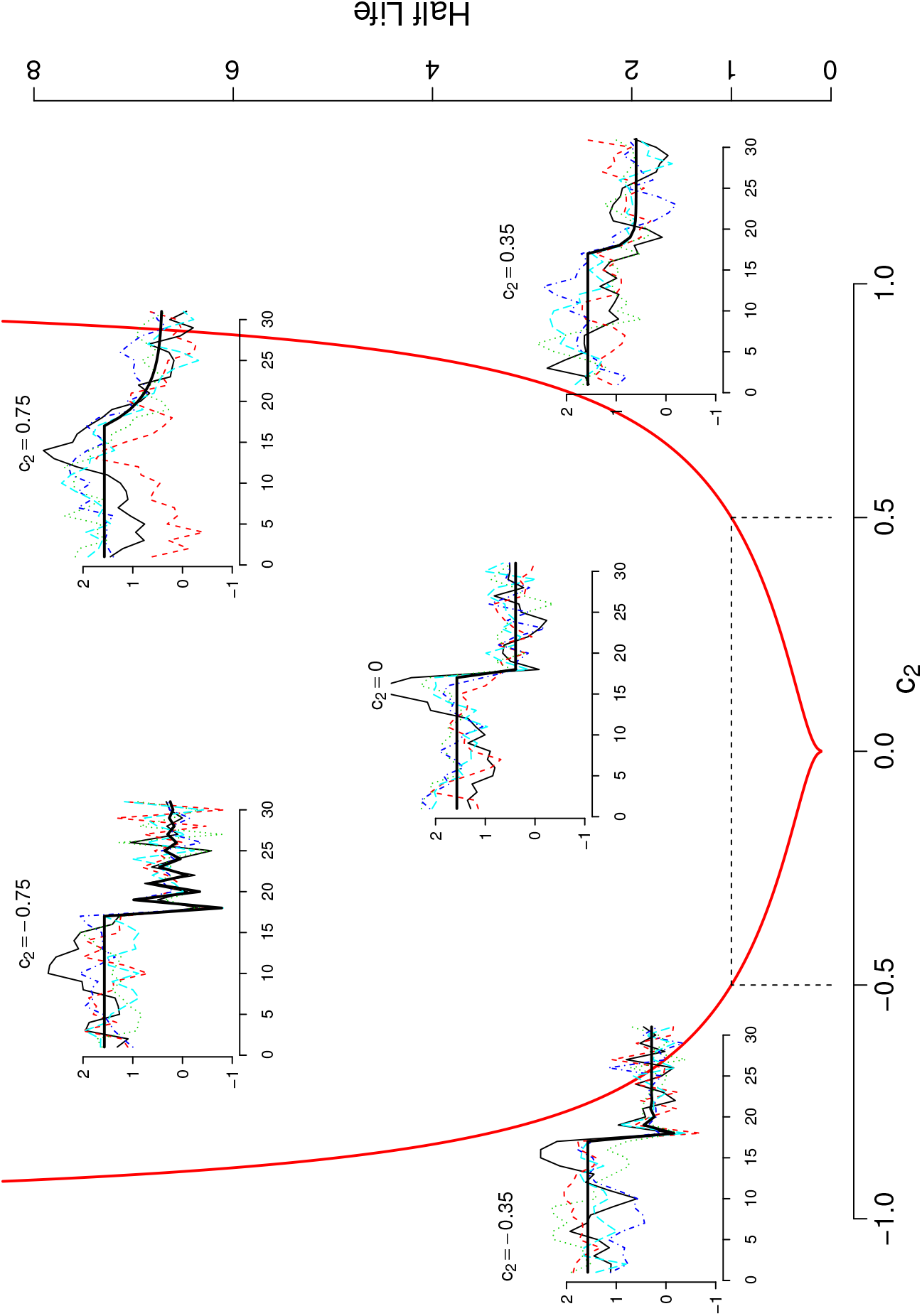
Half life of the change in the mean of the dynamics, 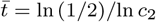, plotted as a function of the strength of density dependence *c*_2_. If the strength of density-dependence after the breakpoint *c*_2_ is close to 1 (*i.e. b* is close to 0 and hence growth is close to density independence), then change occurs slowly. As *c*_2_ approaches 0.5 from the right, the speed of the change increases. When *c*_2_ = 1/2, the change occurs within one time step (*e.g.* a year). If *c*_2_ is exactly 0, then the change is immediate. Between 0 and –1/2 the change still occurs within a single time step, but weak damped oscillations towards the equilibrium are produced. Finally, when –1 < *c*_2_ < –1/2, the speed of the change is slow again, with the added complexity that such change is accompanied with strong, damped oscillations towards the equilibrium. Thus, the non-stationary trend contains information about *a)* the location of the past stationary mean, *b)* the location of the future stationary mean, and *c)* the duration of the transitionary period.

**Result 3:** *The covariance between a population abundance before the breakpoint and after the breakpoint decays geometrically over time in powers of c_2_ to the right of the breakpoint and in powers of c_1_ to the left of the breakpoint. The intensity of such covariance is always proportional to the variance of the stationary distribution of the first partition.*

The partitioned representation of the log-population process X is completed by writing its variance covariance matrix as a block matrix:

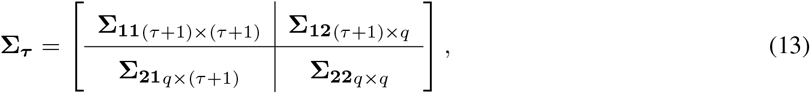

where the diagonal matrices **Σ_*ii*_**, *i* = 1, 2 are the covariance matrices for the pre and post-breakpoint elements of **X** respectively. Accordingly, the sub-matrix **Σ_11_** specifies the variance of the process at every time point before the breakpoint and the covariances among these points. Likewise, the sub-matrix **Σ_22_** specifies the variances at every point after the breakpoint, along with the covariances among these points. As we will see below, such variances and covariances also bear information about the process before the breakpoint. Finally, the sub-matrix **Σ_21_**, which is equal to **Σ_12_′**, specifies the covariance Cov(**X_2_**, **X_1_**), between any time point before the breakpoint and any point after it. Remarkably, these unwieldy expressions reduce to simple mathematical forms that can be easily interpreted. To find these explicit expressions, we first recall that **Σ_*τ*_ = ΓΦΓ′**, and thus that it is convenient to also partition the matrices **Γ** and **Φ** into four sub-matrices matching the dimensions of the Σ_*τ*_ blocks. That is,

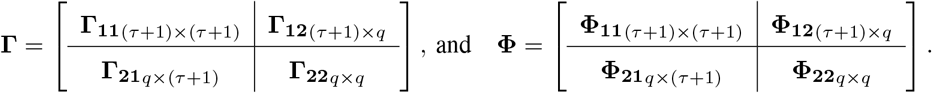

Next, noting that **Γ_12_ = 0**, the variance-covariance of the process can be re-written as

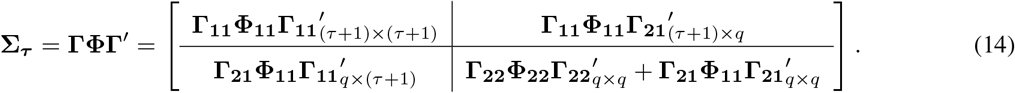

Elementary matrix multiplications and an examination of the power patterns (see Appendix 2) leads to a simple expression of the elements of the matrix **Σ_12_ = Γ_11_Φ_11_Γ_21_′**, which represents the covariance between any process realization before the breakpoint *X*_0_, *X*_1_,…, *X*_τ_, and any process realization after the breakpoint, *X*_τ+1_, *X*_*τ*+2_,…, *X_τ+q_*:

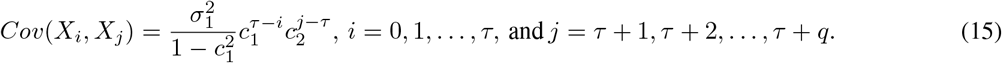

Thus, these covariances are proportional to the variance of the stationary distribution of the first partition and their strength increases from the left of the breakpoint in powers of *c*_1_, and to the right of such point, it decreases in powers of *c*_2_.

**Result 4:** *The variance-covariance matrix **Σ22** can be decomposed into two variance component matrices. An examination of these components reveals exact analytical expressions detailing information about the past and the future stationary distribution of the population process concealed in the post-breakpoint time series.*

While the sub-matrix **Σ_11_** is identical in form to the variance covariance matrix of the process without a breakpoint (eq. A.6, except that with dimensions (*τ* + 1) × (*τ* + 1)), the sub-matrix **Σ_22_** is the sum of two matrices, 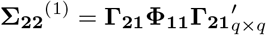 and 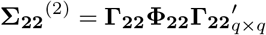.

**Result 4.1** *The first variance component of the matrix **Σ_22_**, given by 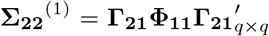, is proportional to the variance of the stationary distribution in the first partition. The weight of such stationary variance decays over time, in powers of *c*_2_.*

After performing the matrix multiplications in the right hand side (RHS) of 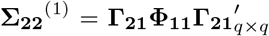 and simplifying the resulting expressions (see Appendix 2) the elements of **Σ_22_^(1)^** are found to be:

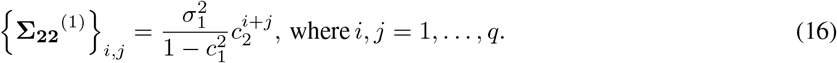

Thus, the first variance component of the second time series partition is proportional to the variance of the stationary distribution in the first partition, 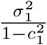. The influence of such stationary variance decays over time, in powers of the second dynamics’ density-dependent effect, *c*_2_.

**Result 4.2** *The second variance component is proportional to the stationary variance of the second partition. These covariances measure the departure of the variance-covariance matrix of the second partition from its limiting form at stationarity. The magnitude of the departure from the limiting covariances is controlled by the strength of density dependence in the second partition, modulated by the positioning of the observations relative to each other and to the location of the breakpoint.*

The elements of the second matrix 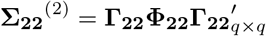 are given by

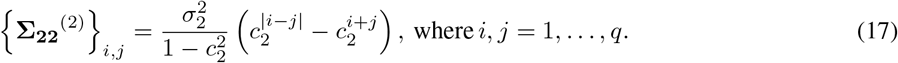

To understand this formula, note that 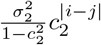 is the covariance between two observations |*i* – *j*| time units apart, had the second partition arisen directly from the stationary distribution of the process parameterized with the second set of parameters. Thus, this second matrix is a measure of the departure of the variance-covariance matrix of the second partition from what the variance-covariance matrix would be if the process, right after the breakpoint, had been drawn from the second stationary distribution. Thus, it follows that the matrix **Σ_22_** can be expressed as a sum of two variance components, one directly related with the stationary variance of the process from the first partition, and another measuring the divergence from stationarity in the second partition in the variance-covariance space. In the particular case when *i = j* (*i.e.* for the diagonal elements of **Σ_22_**), this sum gives the variance of the elements after the second partition. This brings us to our result:

**Result 4.3** *The variances of the log-observations after the breakpoint (the diagonal elements of the variance-covariance matrix **Σ_22_**) can be written as a weighted average of the stationary distribution before and after the breakpoint.*

The developed representation of the transitional variances using the variance decomposition from “Result 4.1 and 4.2” turns out to be analogous to the the equation for the transtional means (eq. 11). Specifically, these variances can be written as a weighted average of the variance of the stationary distribution before and after the breakpoint:

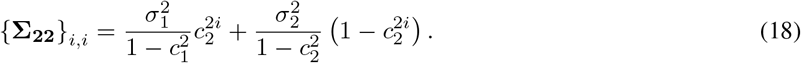

The half life of the change in variance, denoted 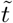 and found by setting

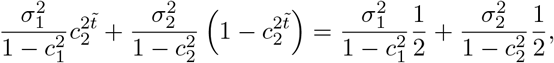

and solving for 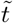 is given by

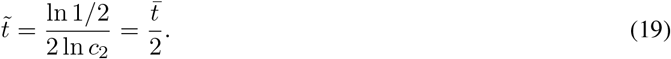

Therefore, the variance of the process transitions twice as fast as the mean of the process from one dynamical scenario to the next. A depiction of these formulae accompanied with matching empirical trends is shown in Fig. 3

**Figure 3:**
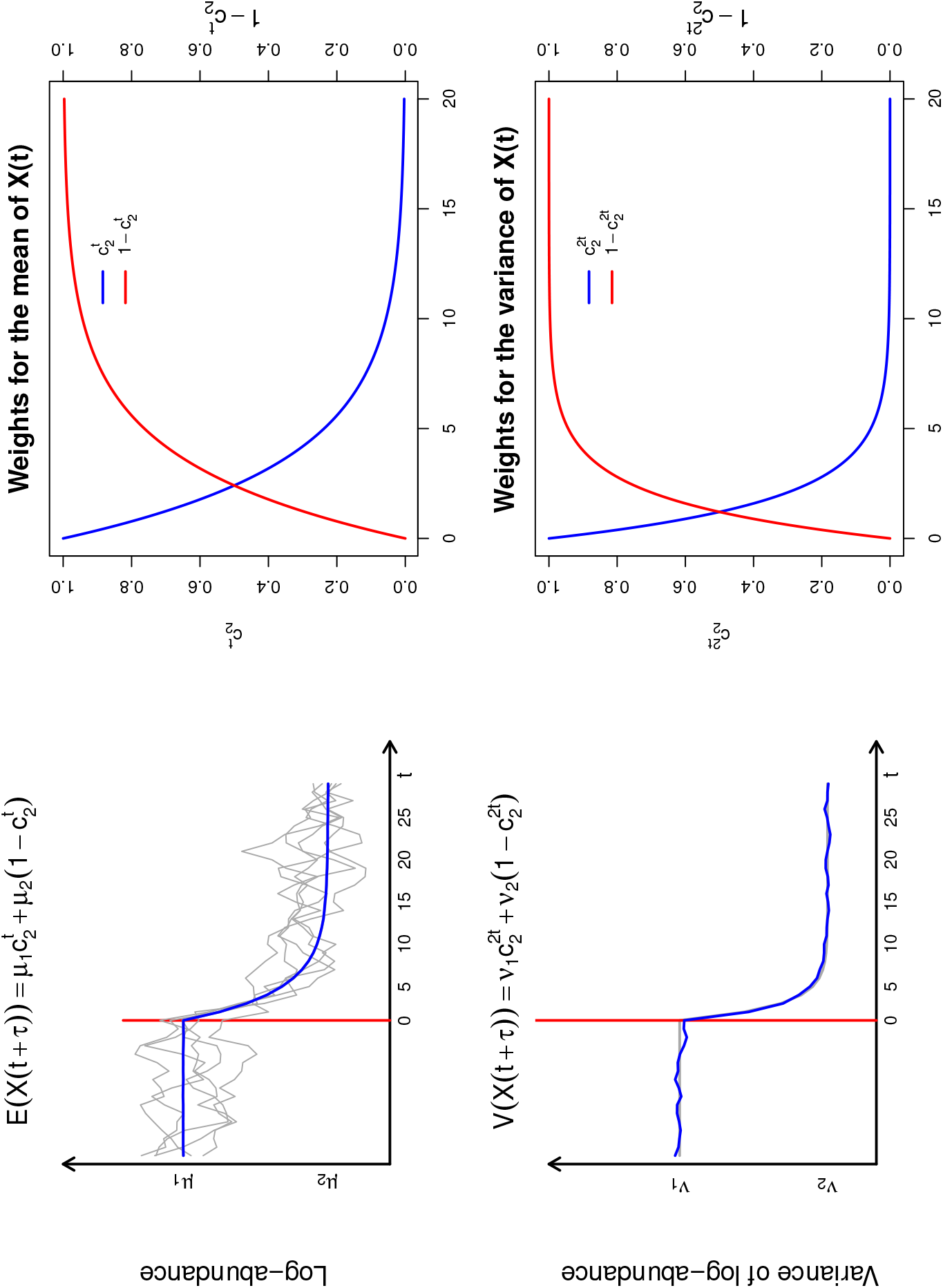
The theoretical and empirical means and variances of the breakpoint Stochastic Gompertz model. In the upper left and lower left panels, the theoretical mean and variance (grey) of the breakpoint process is depicted. Overlaid in the upper panel are 5 simulated trajectories and in blue is the empirical average of 10000 simulated trajectories. The empirical average of the variances of the same 10000 trajectories is depicted in blue in the lower left panel. As stated in the results, both the mean and the variance of the process after a change are a weighted average of the stationary means and variances before and after the change. The upper and lower right panels show how the weights in these weighted averages depend on the strength of density-dependence after the change (See result 1). In particular, as stated in Result 4.3, note that the weights of the variance change twice as fast as the changes in the mean (see for instance [36])

**Result 5:** *The variance-covariance matrix of the observations **Y**, the process with added sampling error, is written as the sum of the variance-covariance of the process and the variance-covariance matrix of the observation error model.*

According to the sampling error setting (see *Parameter Estimation* subsection above),

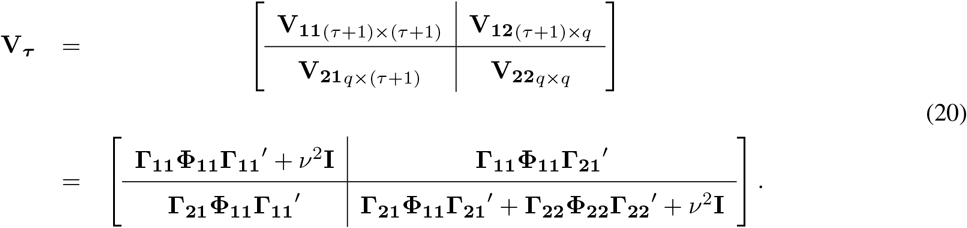

From this development, it follows that the log-likelihood needed for parameter estimation accounting for a change-point in the dynamics occurring between times *τ* and *τ* + 1 is simply the probability density function (pdf) of such MVN distribution evaluated at the observations:

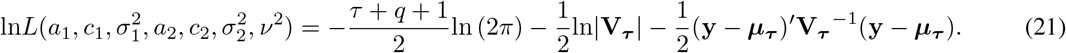

A by-product of this multivariate normal formulation is that the problem of estimation in the presence of missing observations in the time series is readily solved (see Dennis and Ponciano 2014): suppose that a subset *s* of elements of the vector **Y** is missing. Then, the likelihood function of the remaining observations is simply written as in eq. 21 but by removing the *s*^th^ elements in the mean vector ***μ_τ_*** and the *s*^th^ rows and columns of the matrix **V_τ_**.

**Result 6:** *Despite our model being a simplified representation of a change in population dynamics, statistical inferences from this model are robust to biologically realistic departures of the model assumptions.*

After simulating from a model with both, demographic and environmental stochasticities as well as sampling error, the half-life estimates obtained from fitting the BPGSS model appear reliable and with almost no bias, even at the most extreme level of simulated compensatory dynamics (see Fig. 4). The computer code in **R** used to do the simulations, parameter estimation and plotting of each one of the figures is provided in the first author (**JMP**)’s laboratory web page.

**Figure 4:**
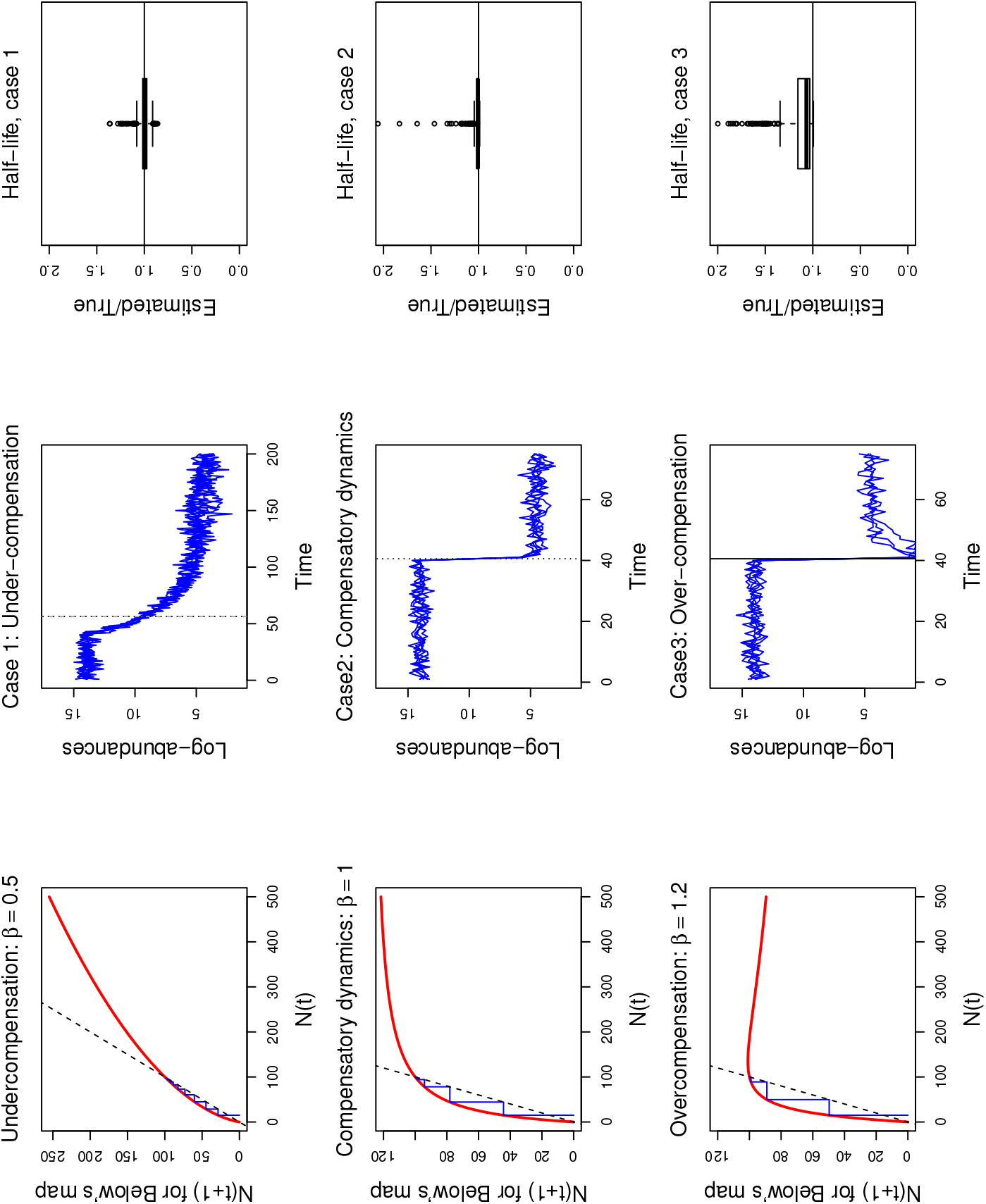
Testing the robustness of the breakpoint Stochastic Gompertz model. A biologically realistic change-point stochastic process was used to test the robustness of the Gompertz model to unaccounted biological complexities. Accordingly, for three different types of compensatory dynamics, 1000 simulations from a population growth model with demographic variability, demographic noise, sampling error and a change-point were used to estimate the parameters of the BPSG model with added sampling error, along with the half life of the change. In the figure, each row, from left to right, shows the type of compensatory dynamics used in the simulations (left panels), 10 samples of the simulated time series of length 200 with the true half life 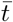 (center panels and solid vertical line) and the estimated half life (dotted vertical line), and a box plot of the 1000 half life estimates (right panels). For the compensatory and over compensatory dynamics, only the first 75 time steps of the simulated time series are shown for clarity.

## 5. Discussion

In this study, we show that after a saltational change in population dynamics’ parameters triggered by a change in environmental conditions, the transition to new equilibrial conditions is governed by the strength of density dependence. In response to a change in environmental conditions, the stationary probability distribution of a species’ population size undergoes a transition and asymptotically approaches a new equilibrium distribution. The statistical properties, means, variances and covariances, respond to change in different ways (Results 1 through 6 above). Remarkably, in all of them it is density dependence what governs the speed of change. In all of these statistical properties, the history of the past persists and the future is anticipated as an explicit function of the strength of density dependence.

According to our second result, the strength of density dependence controls the average of a first passage time, the average time it takes to complete the transition from one stationary mean to the other. This result automatically connects the strength of density dependence with any theoretical dynamic concept. Take for instance the concept of “potential functions”. For continuous time, deterministic population dynamics models of the form dn/dt = m(n), the potential function is defined as the function *u*(*n*) satisfying 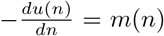 [58]. This function is deeply connected with deterministic waiting times. As Dennis et al [58] point out, the waiting time *t*(*n*) needed to reach abundance n from an initial abundance *x* can be written as a function of the potential function as follows

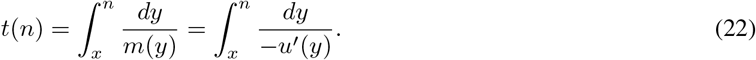

Thus, the formulation in equation 22 aids ecological interpretation because it gives to the potential function a direct connection with *t*(*n*).

The famous “marble in a bowl” conceptual image has been extremely fruitful in Ecology to explain classical dynamic ideas like resilience [59]. The potential function *u*(*n*) is the mathematical representation of the “bowl” in such image. Just as the potential function is connected to deterministic waiting times (equation 22), one could expect our average first passage times, that depend on the strength of density dependence, to be connected with a stochastic formulation of the potential well. In fact, as depicted in ecological textbooks, the “bowl” is often shaken (*e.g.* Figure 6.15 in [59]). This stochastic formulation would then be the mathematical representation of dynamics under such perturbations. Importantly, following our results this representation of a stochastic well will not only depend on the strength of density-dependence but also on the variance properties of the system, which we have concisely described in this paper. We are developing a full treatment of the stochastic potential well in a separate manuscript.

Despite our model being a simplified representation of a change in population dynamics, statistical inferences from this model are robust to biologically realistic departures of the model assumptions. Indeed, here we show that the inference via ML seems to be robust to the inclusion of an atypical yet realistic form of environmental noise: temporal variability in the strength of density dependence, recently shown to be important in animal populations [53] (Fig. 4).This robustness confers the model the generality that is needed for sound hypothesis-driven learning in population dynamics modeling. After simulating from a model with both, demographic and environmental stochasticities as well as sampling error and notably, multiple forms of density-dependence, the half-life estimates obtained from fitting the BPGSS model appear reliable and with almost no bias, even at the most extreme level of simulated compensatory dynamics (see Fig. 4). These results seem to support the early result of [60] who found that the growth rate parameterization of the theta-logistic model that was fairly common across a wide taxonomic spectrum was basically equivalent to the Gompertz model parameterization. Providing a mechanistic support for this pattern is an interesting topic for further research using ecological and evolutionary arguments (see for instance [61]).

Bias in ML estimates of population dynamic models has been previously reported and remedied using REML estimation. In our applied conservation biology work (not shown here) using the GSS model, we have successfully applied the REML bias reduction technique (for an introduction to this topic in population dynamics models see [7]). A full exploration of this bias-reduction issue warrants an independent research study. Therefore, we believe that after sufficient investigation and testing our model may turn out to be sufficient for understanding, management and importantly, prediction. The latter is the focus of our ongoing Population Viability Monitoring (VPM) programatic research [4].

A more complete characterization through simulation of the statistical properties of the estimators of this model is in progress and will be reported elsewhere. In particular, a computational study of the properties of the change point parameter estimate and of the biological parameters as a function of the length of the time series before and after the break are still needed. Given the covariance structure of the model’s second partition *(i.e.* the time series post-change point), we expect the length of the first partition, as well as the size of the change to be critical to be able to correctly estimate the covariance structure in the second partition. In that respect, the location of the change point within the time series is expected to play a critical role. We also expect that these statistical studies will reveal a suitable approximation for the usually less biased restricted maximum likelihood (REML) estimators. In any case, however, if a change point is suspected to be present in a particular data set, extensive statistical tests should be carried to confront our change-point model to a null model of no change, and possibly, to other types of dynamics (see for instance [62]). Interestingly, because the covariance structure of the second partition also contains information about the parameters in the first partition, one could envision estimating the pre-breakpoint parameters even if only post-breakpoint time series data immediately after the breakpoint is available. An immediate application of our results and observations is the study of the recovery process, or the case where the change is from a lower stationary distribution to a (potentially) higher stationary density (see lower panel of Fig. 1). The explicit changes in variances and covariances of the process could then be directly linked to changes in, for instance, pseude-extinction dynamics.

The principles driving the inferences from these models are applicable to a wide array of systems, from macroscopic to microscopic scales. The advent of DNA-based technology, for instance, has made it possible to explore the population dynamics of entire microbial communities, whose signature trends exhibit sudden changes. Being able to achieve a process-based understanding of such changes is at the center of very active research in microbial ecology [63], because different microbial community compositions and structures are associated with different pathological states of the hosts. It follows that understanding the drivers of the “stability” of the community composition is key to understand how to manage changes in a microbial community. Our model can be readily extended to a multi-species autoregressive format (e.g. [36]) that can serve as the basis to move the research from the study of patterns of change in abundance to the study of patterns of change in the nature and intensity of ecological interactions. Such move would make readily accessible an ecological, process-based understanding of change. Much recent work in ecology and other scientific fields aims at developing statistical tests to anticipate changes of state [64]. One of the latest examinations of the problem [65] emphasizes the importance of model-based approaches to find reliable indicators of impending regime shifts. In this paper, we shift the attention from the problem of detecting when a change will occur, to another non-trivial puzzle: understanding and predicting the post-breakpoint behavior of the population dynamics. In this work we show that population time series transitioning from one stationary distribution to another contain explicit information about where the process was before the change-point, where is it heading and how long it will take to transition. We view the current contribution’s explicit modeling as a step in understanding the relationship between stochasticity, density dependence, population dynamics shifts, and the influence of the past dynamics on future dynamics.

This paper has begun the study of the effects on natural population of environmentally induced changes from the simplest manifestation of change −a single saltational break in parameter values. Our experience with many wildlife time-series have led us to believe that, while such sharp changes are not uncommon, they are certainly not universal. This observation raises several questions. How can we investigate the effects of more gradual change? And, how can we distinguish saltational parameter change from more gradual shifts?

Fortunately, the formulations of this paper can be immediately extended to by considering every time step as a potential saltational change as in [66]. As full likelihood estimates will be available for any modelable transition of parameter values, identifying the best model of change can be accomplished using information criteria. This also is material for upcoming research.

## Acknowledgements

JMP and MLT were funded by both NIH R01-GM103604 and by a Montana Fish, Wildlife, and Parks contract 060327 to M. L. Taper.

## AppendixA

## AppendixA.1. Matrix formulation of the breakpoint process

Here we derive by induction a matrix formulation for the (random) discrete map with a breakpoint. To do that, we derive first the multivariate distribution of the process without a breakpoint, initialized at time 0 and stopped at time *q*. Although this derivation has been shown before (Dennis et al 2006), presenting it here illuminates the formulation of the multivariate distribution with a change-point. To find the joint distribution of [*X*_0_, *X*_1_, *X*_2_,… *X_q_*]′ we first iterate the stochastic Gompertz map in eq. (4):

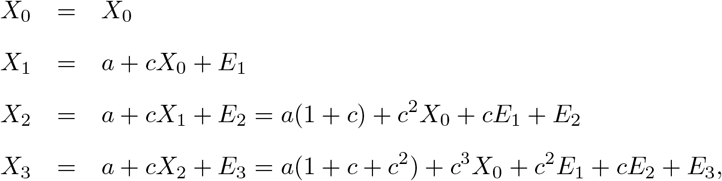

Then, continuing in this fashion and using vector notation we get

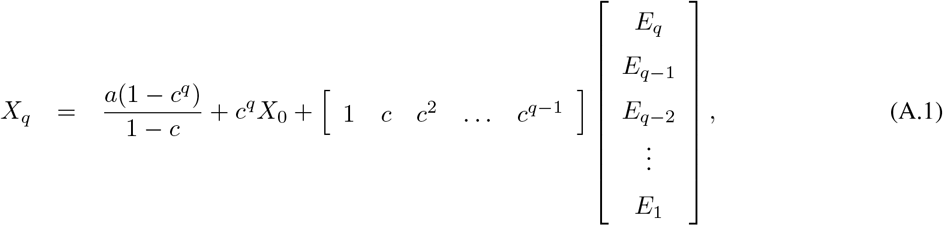

where we used the well known formula for the partial sum of a geometric series. Upon examination of this iteration, it is seen that the vector of log population sizes through time, **X** = [*X*_0_ *X*_1_ *X*_2_ … *X_q_*]′ can be written as:

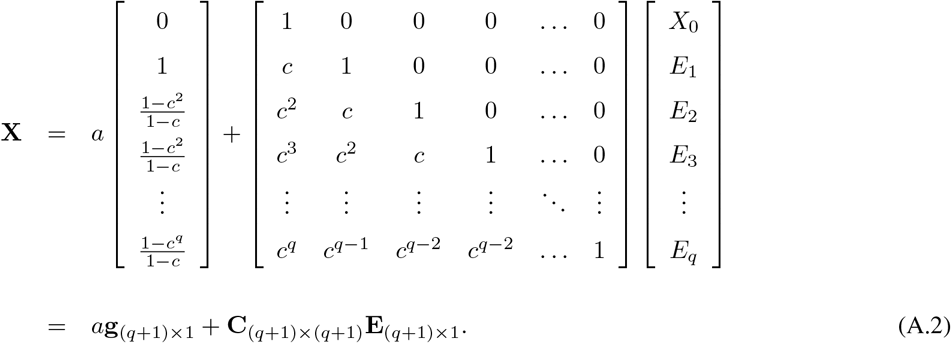

Assuming that the first observation arises from the Gaussian stationary distribution with mean and variance *a*/(1 – *c*) and *σ*^2^/(1 – *c*^2^) respectively, we then have that:

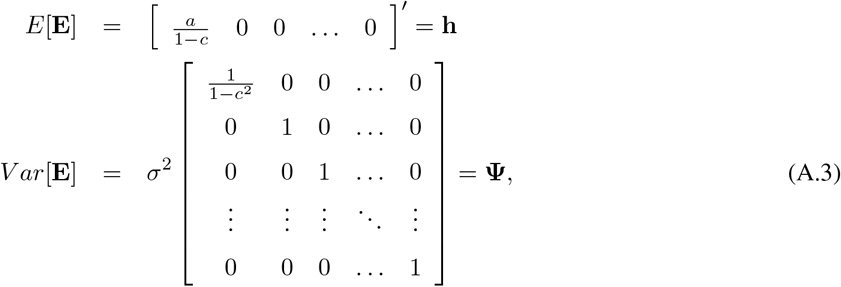

and

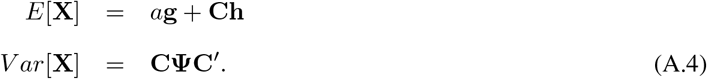

Using this multivariate mean and variance and the properties of the Multivariate Normal (*MVN*) distribution (Johnson and Wichern 2002, chapter 4), it follows that

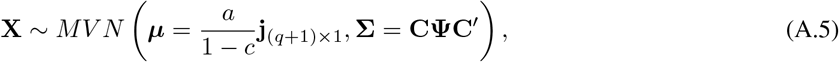

where

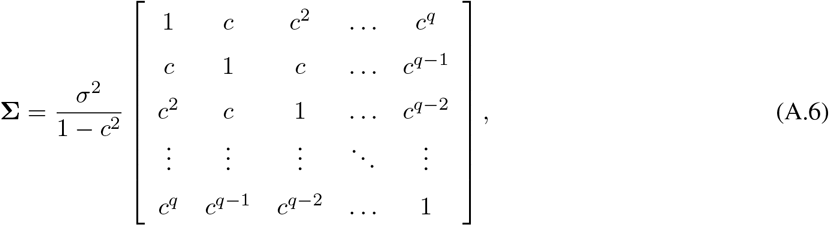

and **j**_(*q*+1)×1_ is a vector of ones of size *q* + 1.

Now, to derive the multivariate distribution of the process with a breakpoint, we start by writing expressions for the log-population process, from time *τ* (right before the breakpoint) and onwards:

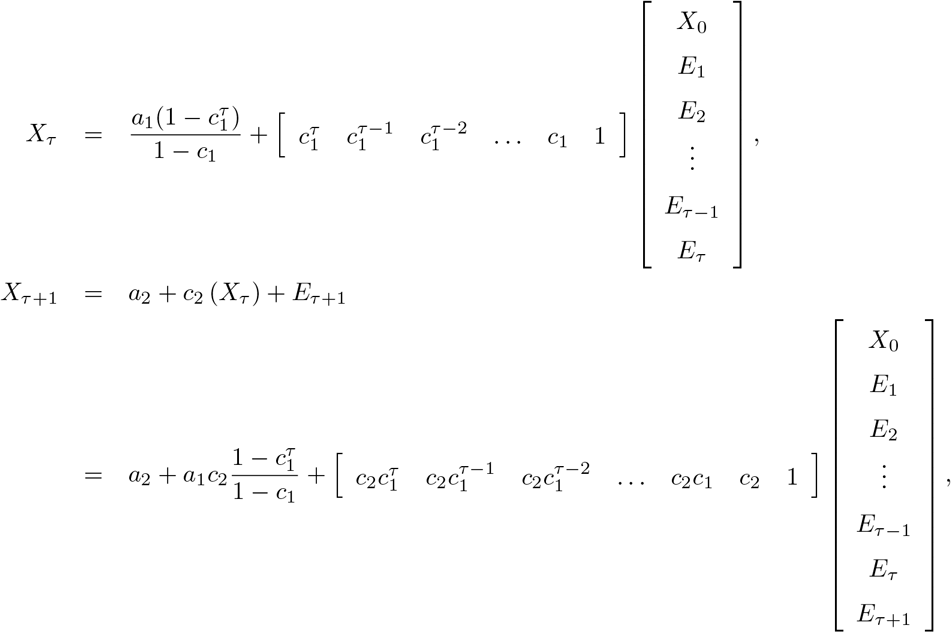

and continuing in this fashion, a geometric series for *c*_2_ appears in the solution:

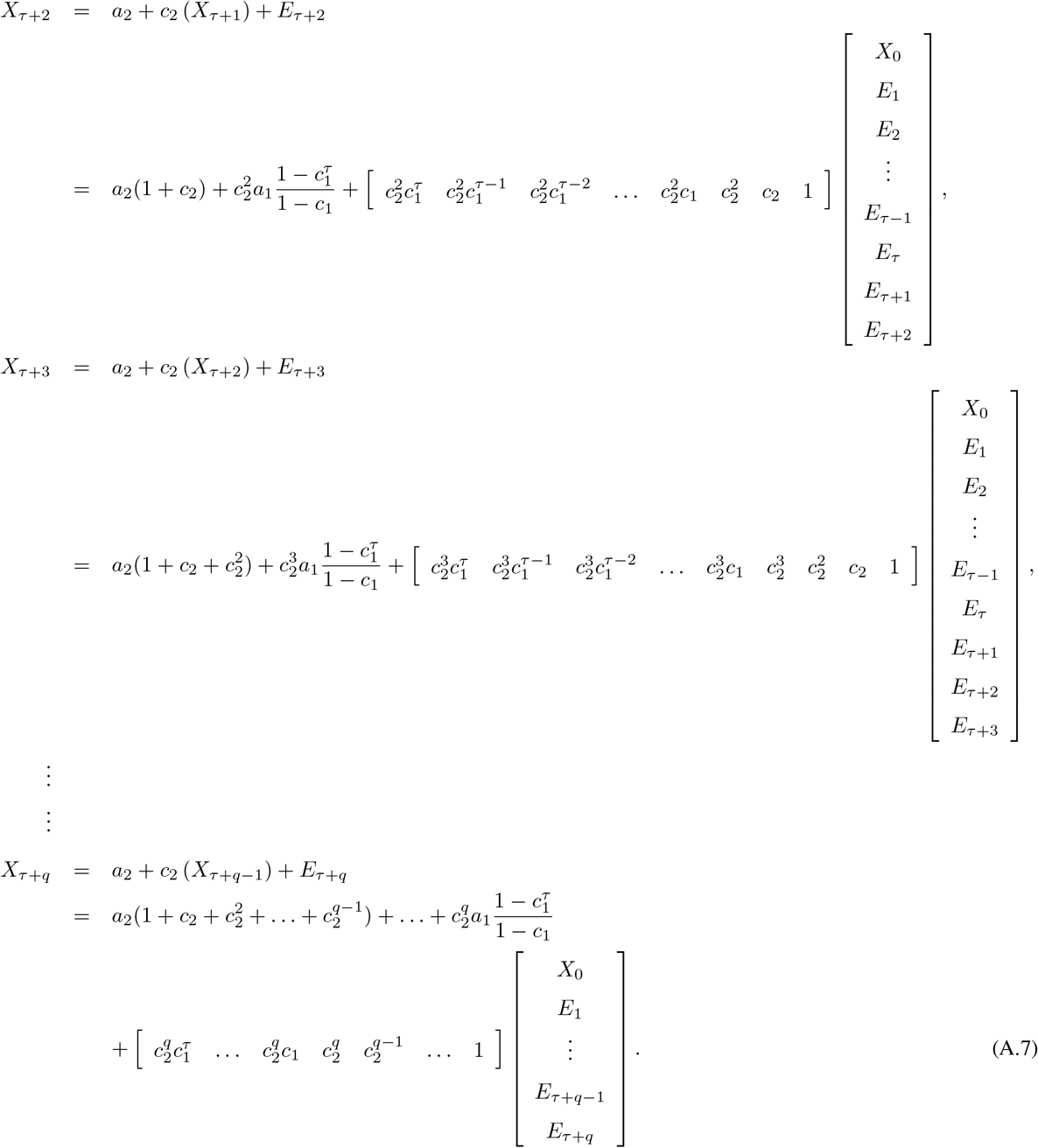

The induction patterns revealed by this iterated map can then be readily expressed in the matrix form shown in eq. (7) in the main text.

## AppendixA.2. Derivation of the variance-covariance matrix of the breakpoint process

Here we derive the four sub-matrices of **Σ_*τ*_**. We recall that we also partitioned the matrices **Γ** and **Φ** into four sub-matrices matching the dimensions of the **Σ_*τ*_** blocks. For Γ these matrices are

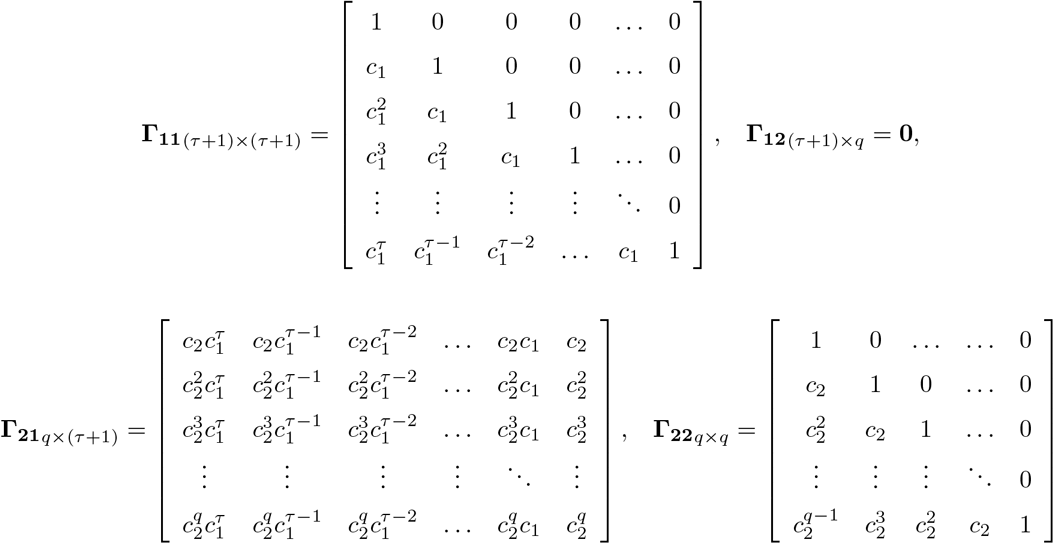

and for **Φ** these are

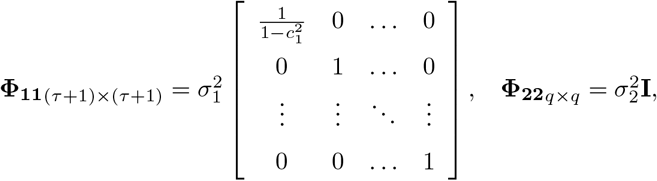

and

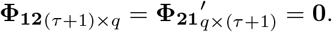

With these matrices in place, the matrices in eqs. (16) and (17) are obtained by simple matrix multiplication and elementary algebraic manipulations:

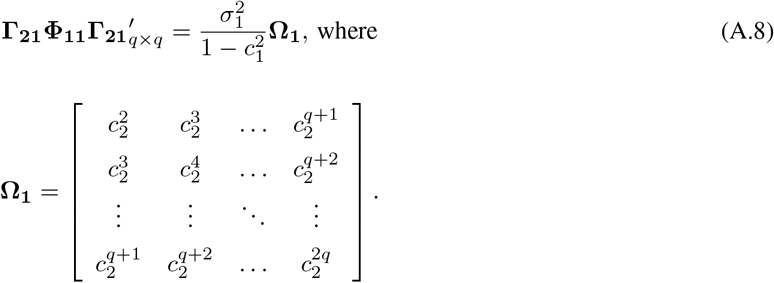

whereas

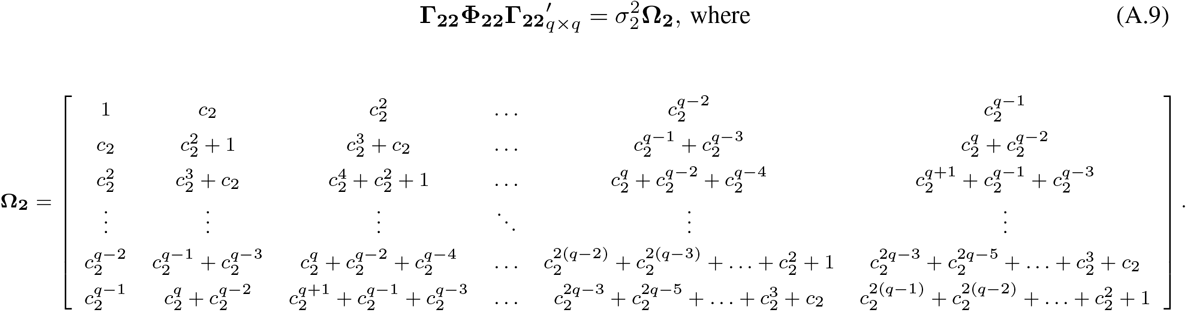

Noting that

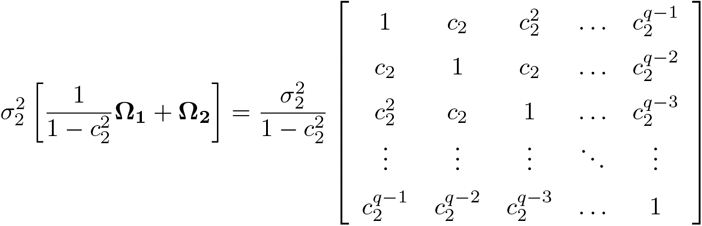

is identical to the variance covariance of the second partition, had it arisen directly from the stationary distribution of the second set of parameters, and solving in that equation for 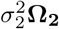 leads to the expression in eq. (17).

Next, we compute **Σ_12_** = *Cov*(**X_1_, X_2_**), which is equal to the transpose of Σ_21_. Straightforward matrix multiplications yield

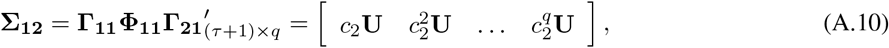

where

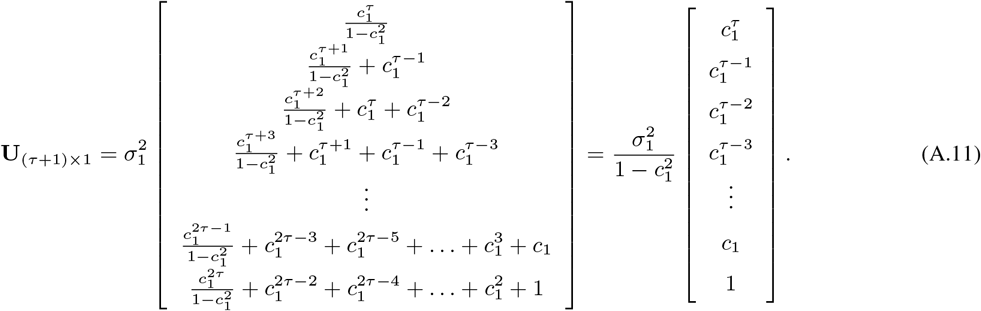

A simple examination of eq. (A.10) and eq. (A.11) then leads to eq. 15 in the main text.

